# Primer- and template-independent RNA polymerization by terminal nucleotidyltransferase TENT4B

**DOI:** 10.64898/2026.03.05.709691

**Authors:** Neha Nagpal, Albert K. Tai, Yick W. Fong, Suneet Agarwal

## Abstract

RNA synthesis by eukaryotic polymerases requires existing polynucleotides to serve as templates or primers. Here, we describe primer- and template-free RNA generation by human terminal nucleotidyltransferase 4B (TENT4B) via de novo polymerization of free nucleotides. We observed that recombinant TENT4B (rTENT4B) consumes ATP to yield inorganic pyrophosphate in the absence of a primer or template, concurrent with the appearance of oligomeric poly-adenosine RNA products. Remarkably, 5’ labels on γ-phosphate-modified ATP or GTP are retained during polymerization in the presence of unlabeled nucleotide triphosphates (NTPs). These polymers are created at a similar efficiency irrespective of the inclusion of a primer, indicating robust RNA synthesis by rTENT4B from free NTPs. While canonical purine NTPs are favored, nucleotide diphosphates can also serve as substrates for rTENT4B-mediated de novo RNA polymerization. rTENT4B-mediated RNA synthesis using free adenosine nucleotides shows high processivity to generate 1000s-mers, whereas guanosine nucleotide polymerization is strongly and uniformly self-limited and yields a 3’-exonuclease-resistant oligonucleotide. Interrogation of other RNA polymerases reveals potential capacity for de novo polymerization using free ATP, albeit at significantly higher substrate concentrations and lower efficiency compared to rTENT4B. Our data provide definitive evidence of efficient template-free *de novo* RNA synthesis by a eukaryotic polymerase.

## Introduction

RNA polymerases use nucleotide triphosphates (NTPs) to generate polymers using pre-existing nucleic acids in two non-mutually exclusive ways: (1) template-dependent polymerization to create a product complementary to a DNA or RNA sequence, and (2) primer-dependent polymerization via 3’ extension of another RNA species. Template-dependent RNA polymers are mainly formed *de novo* starting with two free NTPs by mechanisms well-characterized at the biochemical and structural levels for diverse polymerases ranging from eukaryotic RNA polymerase II to T7 bacteriophage RNA polymerase, which is widely exploited in biotechnology(1-4). In most cases, template-dependent polymers begin with a free purine NTP (ATP or GTP) opposite a templating pyrimidine with synthesis of a short RNA, followed by extensive coordinated structural changes of the transcription complex into an efficient template-directed elongation machinery(5). Template-directed RNA polymerization by viruses can also be initiated from the free hydroxyl group of a pre-existing nucleic acid or protein(6,7), but generally, primer-dependent RNA polymerization refers to template-independent polynucleotide addition to existing short or long coding and non-coding RNAs (ncRNA) by ribonucleotidyl transferases(8). Amongst the best characterized of these are canonical poly(A) polymerases (PAPs) that modify mRNAs in a termination-coupled manner downstream of endonucleolytic cleavage at the AAUAAA sequence motif, thus promoting mRNA stability, nuclear export, and translation(9,10). The CCA-adding enzyme is a notable ncRNA-dependent RNA polymerase that undergoes structural changes to ensure addition of a precise, untemplated trinucleotide to achieve functional tRNA maturation(11). Despite these diverse properties, there are no examples of known polymerases using neither a primer nor template for RNA synthesis.

Distinct from canonical PAPs, noncanonical template-independent RNA polymerases in eukaryotes are grouped under the category of terminal nucleotidyl transferases (TENTs) and serve varying RNA quality control and maturation functions(8,12,13). Eleven TENTs have been identified in humans, sharing homology in the nucleotidyltransferase domains with PAPs and each other, but with varying NTP and RNA substrate preferences(14). These include the terminal uridyltransferase TUT1 (aka TENT1) which modifies U6 snRNA to promote its biogenesis, and TUT4 (aka TENT3A) and TUT7 (aka TENT3B) which oligouridylate microRNAs (miRNAs) to alter their inhibitory function and cause RNA degradation(15,16). A similar role in miRNA regulation is played by TENT2 but via oligoadenylation(17). MTPAP (aka TENT6) encodes the mitochondrial PAP which oligoadenylates and regulates the function of mitochondrial mRNAs and tRNAs(18,19). The roles of the four TENT5 family members are emerging as diverse and include innate immunity, gametogenesis, and secretory functions in different cell types (20-23). The remaining family members TENT4A and TENT4B are unusual amongst TENTs in displaying a relaxed substrate specificity, primarily utilizing ATP but also GTP, followed by CTP and UTP(24-28). This promiscuity may be relevant in vivo for mixed adenylyl and guanylyl tailing in the stabilization of mRNAs(29). Interestingly, TENT4A and TENT4B have been found to play critical roles in polynucleotide tailing of RNAs from viruses including hepatitis B virus (HBV) and hepatitis A virus (HAV)(30-34). Distinct roles for TENT4B have also been shown in oligoadenylation of several human non-coding RNAs including telomerase RNA (TERC), small nucleolar RNAs (snoRNAs), ribosomal RNAs, and miRNAs(26,35-40).

TENT4B has emerged as a therapeutic target due to its role in the regulation of nascent TERC and HBV surface antigen (HbSAg) RNA. A phenotypic screen to reduce HbSAg expression from hepatocytes, a hallmark of chronic infection, yielded potent small molecule dual inhibitors of TENT4A and TENT4B, revealing their unexpected role in viral mRNA formation and thus providing new targets for chronic HBV infection (31,41,42). Independently, we and others identified a role of TENT4B in destabilizing TERC, the non-coding template for telomerase that maintains telomere length and stem cell self-renewal capacity(24,35,43-45). TENT4B oligoadenylates and targets nascent TERC transcripts for degradation by the nuclear exosome, a process that is critically opposed by the exonuclease poly(A) ribonuclease PARN(26,35,36,46,47). In genetic forms of human telomere biology disorders (TBDs), *PARN* loss-of-function leads to unopposed TERC destabilization by TENT4B, and in turn reduced TERC RNA, telomerase insufficiency, and myriad diseases due to premature stem cell exhaustion. These findings led us to screen for small molecule inhibitors of recombinant TENT4B (rTENT4B) as a potential pharmacological approach to restore telomerase function in individuals with genetic TBDs(24). Using TENT4B inhibitors from our screen as well as those identified in HbSAg studies, we showed restoration of TERC RNA steady-state levels and telomere length in vitro in patient-derived induced pluripotent stem cells and in vivo in human hematopoietic stem cells in a mouse xenotransplantation model of TBD(24,30,42).

While performing the high throughput inhibitor screen, we unexpectedly identified novel activities of rTENT4B in RNA synthesis. Here, we demonstrate that rTENT4B can initiate RNA polymerization from free ATP or GTP in the absence of a primer or template. rTENT4B shows highly processive de novo RNA polymer formation from free NTPs, similar to using an oligonucleotide primer. We find that purine nucleotide diphosphates can also be used as substrates for de novo chain synthesis by rTENT4B, albeit at lower efficiency. Targeted evaluation of other canonical and noncanonical polymerases shows their variable capacity for primer-and template-independent RNA synthesis, but at higher ATP concentrations than rTENT4B and without definitive proof of primer-free initiation. Our data defines novel primer- and template-free RNA synthesis capabilities of a eukaryotic RNA polymerase. These results promise to broaden our understanding of RNA polymerase function and evolution and may serve useful in biotechnology applications.

## Materials & Methods

### Purification of recombinant proteins

For recombinant TENT4B (full length, WT), cDNA (Genbank CCB84642.1) was cloned into a modified pMtac-His6 vector and purified as described(24).

For recombinant TENT4B (full length-catalytic site mutant), cDNA (Genbank CCB84642.1) was cloned into a modified pMtac-His6 vector and two aspartate residues in the catalytic domain were exchanged to alanine(25), and the resulting construct was purified as for WT full length TENT4B.

For recombinant TENT2 (full length) purification, codon-optimized TENT2 (NCBI NP_001107865.1) sequence was introduced into a modified pMtac-His6 vector and the purification was performed as described(24).

For recombinant PARN (full length) purification, codon-optimized PARN (NCBI_NP_002573.1) sequence was introduced in a modified pMtac-His6 vector, in frame with a C-terminal FLAG tag, and purification was performed as described(24).

For recombinant TENT4B C-terminal truncation construct (TENT4B-delC) encoding amino acids (a.a.) 1-511 (deletion of a.a. 511-631) of Genbank CCB84642.1, cDNA was cloned into a modified pMtac-His6 vector, expressed and purified by Ni-NTA affinity chromatography as described(24), except that proteins were eluted in elution buffer with 0.1 M NaCl. Peak Ni-NTA fractions were pooled and applied to Q Sepharose Fast Flow pre-equilibrated in elution buffer (50mM HEPES-KOH, pH7.6, 0.1 M NaCl, 2mM MgCl2, 0.05% NP-40 and 10% glycerol) containing 2 mM DTT, 1 mM benzamidine, 0.5 mM PMSF). Q Sepharose flow-through fraction contains active TENT4B-delC and was dialyzed against elution buffer containing 0.4 M NaCl.

For recombinant TENT4B and TENT4A catalytic cores (for TENT4B- a.a. 115-453 of Genbank CCB84642.1, for TENT4A- a.a. 225-560 of NCBI_NP_008930.2) cDNA was cloned into a modified pMtac-His6 vector, expressed and purified by Ni-NTA affinity chromatography as described(24), except that proteins were eluted in elution buffer with 0.125 M NaCl. Peak Ni-NTA fractions were pooled and applied to Q Sepharose Fast Flow pre-equilibrated in elution buffer (50mM HEPES-KOH, pH7.6, 0.125 M NaCl, 2mM MgCl2, 0.05% NP-40 and 10% glycerol) containing 2 mM DTT, 1 mM benzamidine, 0.5 mM PMSF). Q Sepharose flow-through fraction contains active TENT4B-catalytic core and was dialyzed against elution buffer containing 0.4 M NaCl.

### Luciferase assay

ATP consumption was measured using Kinase-Glo Max reagent (Promega, V6074). Enzymatic reactions with rTENT4B and ATP were performed in a buffer containing 25 mM Tris-HCL, 5 mM MgCl2, and 50 mM KCl, in the presence or absence of 1pmole an RNA oligo (oligo A15) as substrate, in 10µl in 384-well format (Corning, 3820). After 1hr (or at indicated times in the figure) reactions were stopped with 5µl of Kinase-Glo reagent followed by centrifugation at 2000rpm for 1 min. Luciferase signal was read using Envision plate reader (Perkin Elmer). Each reaction was performed in n=3 biological replicates.

### RNA oligonucleotides

RNA oligonucleotides were synthesized by IDT and purified by RNase-Free HPLC.

Oligo1- /5’-6-FAM/rCrUrGrCrCrUrGrCrCrUrGrCrCrUrGrCrCrUrGrC

Oligo2- rCrUrGrCrCrUrGrCrCrUrGrCrCrUrGrCrCrUrGrC

Oligo A15- rArArArArArArArArArArArArArArA

Oligo3- /5’-6-FAM/rCrUrGrCrCrUrGrCrCrUrGrCrCrUrGrArArArArA

### Nucleoside triphosphates

ATP- Adenosine-5’-triphosphate (N0450, New England Biolabs)

ADP- Adenosine-5’-diphosphate (NU-1198, Jena Bioscience)

AMP- Adenosine-5’-monophosphate (A1752, Sigma)

GTP- Guanosine-5’-triphosphate (N0450, New England Biolabs)

GDP- Guanosine-5’-diphosphate (G7127, Sigma)

CTP- Cytidine-5’-triphosphate (N0450, New England Biolabs)

CDP- Cytidine-5’-diphosphate (C9755, Sigma)

UTP- Uridine-5’-triphosphate (N0450, New England Biolabs)

UDP- Uridine-5’-diphosphate (94330, Sigma)

Cy5-ATP- γ-(6-Aminohexyl)-ATP-Cy5- (NU-833-CY5, Jena Bioscience)

Cy5-GTP- γ-(6-Aminohexyl)-GTP-Cy5- (NU-834-CY5, Jena Bioscience)

### Polymerase reactions

For RNA substrate extension assays, reactions were performed using various NTPs or NDPs with rTENT4B or other TENTs. Briefly, 2.5 pmoles of rTENT4B (or TENTs) were added per 10µl reaction containing 1 pmol of RNA oligo1 and 25mM Tris-HCl (pH7.4), 50mM KCl, 5mM MgCl2, plus 100µM ATP or 1mM NDP/ NTP (or indicated concentrations, in figures). Reactions were incubated at room temperature for 1hr (or at time indicated in the figures) and stopped using formamide loading buffer (10mM EDTA and 83.3% formamide) and resolved using denaturing polyacrylamide gels (15% Criterion™ TBE-Urea Polyacrylamide Gel, 26 well, 15 µl, Bio-Rad, 3450093). Gels were imaged using FAM channel of the GE Amersham Typhoon biomolecular imager.

For de novo synthesis or no-substrate reactions, the reactions were performed as above but without an RNA oligo. The reactions products were resolved using denaturing gels as above and synthesized RNA products were detected using Sybr Gold Nucleic Acid stain (ThermoFisher Scientific, S11494), and gels were imaged using Cy3 channel of the imager.

For reactions with CID1 or Poly (U) polymerase (New England Biolabs, M0337S), reactions were performed in enzyme-specific buffer and 1U of enzyme was added per 10µl of the reaction mix. Reactions were incubated at 37°C for 1hr and stopped using Formamide loading buffer and resolved and imaged as above.

For reactions with fluorescent or Cy5-labeled NTPs, reactions were performed using similar buffer conditions as above. Briefly, 250 µM of ATP (or at indicated concentrations, in the figures) or other NTPs or NDPs (at indicated concentrations, in figures) were incubated with 100µM of Cy5-ATP or Cy5-GTP and 2.5 pmoles of rTENT4B or other TENTs. Reactions were performed at room temperature for 30min (or at times indicated in figures) and stopped using formamide loading buffer. Reaction products were column purified (Oligo Clean and Concentrator, D4060) to remove unincorporated nucleotides or fluorescent dyes and resolved using denaturing polyacrylamide gels (15%). Gels were imaged using Cy5 channel of the imager. Post-imaging, to visualize non-fluorescent nucleic acid products, the gels were stained using Sybr Gold and imaged using Cy3 channel of the imager. Reactions with yeast polyA-polymerase or YPAP (Jena Biosciences, RNT-006) were performed using manufacturer’s buffer. Briefly, 300U of YPAP were added per 10µl reaction containing 100µM of Cy5-ATP as above, and 1000µM of ATP (unlabeled) and incubated at 37°C for 30min. Reactions were stopped using formamide loading buffer, and column-purified and resolved as described above. Post-imaging, to visualize non-fluorescent RNA products, the gels were stained using Sybr Gold and imaged using Cy3 channel of the imager.

### Deep sequencing of RNA oligo 3′ extended products

#### Oligo extension

For generation of 3’ extended RNA oligos, reactions were performed using 2.5 pmoles of rTENT4B containing 1 pmol of RNA oligo1 and 25mM Tris-HCl (pH7.4), 50mM KCl, 5mM MgCl2, along with 1µM ATP and 333 µM each of GTP, CTP and UTP. Reactions were incubated at room temperature for 20 min to allow for limited extensions and stopped using formamide loading buffer (10mM EDTA and 83.3% formamide), followed by purification using Oligo clean and concentrator kit (Zymo Research, D4060) and purified products were eluted in RNase free water.

#### Linker ligation and 3’ rapid amplification of cDNA ends (RACE)

The RACE protocol was performed as described(24). Briefly, 100ng of purified reaction products from above were ligated to 5’-adenylated-3’-blocked adaptor (Universal miRNA cloning linker, New England Biolabs; S1315S) using T4 RNA ligase truncated KQ (New England Biolabs M0373S) and PEG 8000. Ligated RNA was then purified, followed by cDNA synthesis and PCR amplification (forward primer-CUGCCUGCCUGCCUGCCUGCA, reverse primer-CTACGTAACGATTGATGGTGCCTACAG).

#### Deep sequencing

Linker ligated cDNAs from above were purified using Qiaquick PCR purification kit (Qiagen, 28106) and used for library preparation using the TruSeq Nano DNA LT Library Prep Kit (Illumina, 20015965). Library preparation was done as per kit protocol. Briefly, linkers carrying unique barcodes were ligated to individual RACE amplicons using DNA ligase followed by amplification with Illumina adapters, and size selection using sample purification beads. The completed libraries were submitted to the Tufts University Genomics Core to perform MiSeq (Illumina platform) with paired-end 250 bases using the Illumina TruSeq V2 500 cycles kit after confirming the quality and quantity of each library sample.

For analysis, sequencing and ligation adaptors were removed from raw reads using three sequential rounds of Cutadapt. Reads with no sequence after adaptors trimming were filtered out. The resulting files were then used as an input for the custom Perl script (counttail.pI) to quantify the distribution of 4 bases by position. The outputs were then normalized to 100% and cumulative proportion of each base (A or C or G or U) for a given extended position was calculated as a fraction of total extended oligo species, and graphed. The script used for analysis is given at https://github.com/alberttai/tailcount/

### RG7834 treatment

For RG7834’s effects on de novo synthesis, 100µM (final concentration) of RG7834 (Medkoo Biosciences, 563793) solution in DMSO was added to 10 µl reaction mix containing 100µM of Cy5-ATP with 2.5 pmoles of rTENT4B in the presence/absence of 100µM ATP, in the absence of an RNA-substrate. Reactions were performed at room temperature for 30min and stopped using formamide loading buffer. Reaction products were column purified (Oligo Clean and Concentrator, D4060) to remove unincorporated nucleotides or fluorescent dyes and resolved using denaturing polyacrylamide gels (15%). Gels were imaged using Cy5 channel of the imager.

### Pyrophosphate (PPi) analysis

To detect the pyrophosphate released during the polymerase reactions, reactions were performed in 96-well plates (Costar 96-well Flat-Bottom EIA plate, BioRad, 2240096). 100-1000µM ATP (or at concentrations indicated in the figures) or 1000µM GTP and 50 pmoles of rTENT4B were incubated in a 100µl reaction containing buffer, dye and kit components (EnzChek Pyrophosphate Assay kit, Thermofisher Scientific, E6645), with or without 25 pmoles of oligo2. Reactions were performed at room temperature for 30min (or at times indicated in the figures), and absorbance was measured at 360nm using CLARIOstar BMG Labtech microplate reader.

For reactions with YPAP and *E. coli.* polyA-polymerase (EPAP), 3000U of YPAP or 25U of EPAP were added per 100µl reaction buffer and other kit components (EnzChek Pyrophosphate Assay kit), and reactions were incubated at 37°C and absorbance was measured as above.

### Differential scanning fluorimetry

Direct binding of NTPs to rTENT4B was detected using an indicator dye SYPRO orange (ThermoFisher Scientific, S6651) diluted 1:5000 in 20µl of buffer containing 20 µM rTENT4B alone or either with 1pmole RNA oligo A15, or with 100 µM NTP, 25 mM Tris-HCL, 5 mM MgCl2, 50 mM KCl. Dye-buffer-protein mixtures were heated from 10 to 95 °C at a rate of 1 °C/min and fluorescence signals were monitored by 7500 Fast Real-Time PCR Systems (Applied Biosystems). Each curve represents an average of three measurements and Thermal Shift software (ThermoFisher Scientific, 4466038) was used for raw data analysis, and GraphPad Prism was used for plotting the melt curves.

### Ribonuclease treatment

*For polymerase reactions using GTP and FAM-labeled RNA oligo:* reactions were performed using various 1000µM GTP with 2.5 pmoles of rTENT4B and 1 pmol of RNA oligo 3. Reactions were incubated at room temperature for 1hr and stopped using formamide loading buffer. Reaction products were column purified (Oligo Clean and Concentrator, D4060) to remove unincorporated nucleotides and reactions products were subjected to exonucleases treatments as below.

For rPARN, 0.001 pmoles of rPARN was incubated with purified reaction products in the buffer containing 25mM Tris-HCl (pH7.4), 50mM KCl, 5mM MgCl_2_, 0.1mg/ml BSA, 0.02% NP-40, 0.01 mM EDTA and 1mM DTT in a 10µl reaction. Reactions were incubated at room temperature for 20 min.

For exonuclease T (New England Biolabs, M0265), 5U of enzyme was used in enzyme specific buffer in a 10µl reaction and incubated at room temperature for 30min.

Reactions were stopped using formamide loading buffer and resolved using 20% TBE-urea polyacrylamide gels. Gels were imaged using a GE Amersham Typhoon biomolecular imager in FAM channel.

*For polymerase reactions using Cy5-labeled NTPs:* reactions were performed using 100µM of Cy5-ATP and 10µM ATP or 1000µM GTP with 5pmoles of rTENT4B (catalytic core), without any RNA oligo. Reactions were incubated at room temperature for 30min and stopped using formamide loading buffer. Reaction products were column purified (Oligo Clean and Concentrator, D4060) to remove unincorporated nucleotides and fluorescent dye components.

Polymerase reactions products were subjected to exonucleases treatments with rPARN and Exonuclease T as described above. For RNase R (Lucigen, NC1163864) treatments, polymerase reaction products were treated with 10U of RNase R enzyme with enzyme specific buffer in a 10µl reaction at 37°C for 10 min. Both exo- and ribo-nucleolytic reactions were stopped using formamide loading buffer and resolved using 20% TBE-urea polyacrylamide gels and imaged in Cy5 channel of the imager.

### Quantification and statistical analysis

Error bars presented mean with standard error. P values were calculated based on Student’s t test or ANOVA (see Figure legends) and p<0.05 was defined as significant. Statistical analyses were done using GraphPad Prism 10 (V10.6.1) software.

## Results

### Primer-independent ATP polymerization by rTENT4B

During our high-throughput screen to identify TENT4B inhibitors, we observed that ATP was consumed by rTENT4B in the absence of an RNA oligonucleotide substrate. Specifically, our screen measured residual ATP substrate in an rTENT4B primer extension reaction via luciferase-mediated light intensity, an elevation in which would indicate enzyme inhibition. Interestingly, we observed a similar ATP concentration-dependent decrease in luciferase signal over time irrespective of oligonucleotide addition (**Fig. 1A**). To determine the basis of the loss of luciferase signal, we assessed whether ATP hydrolysis products pyrophosphate (PPi) versus inorganic phosphate (Pi) were being generated, as measured using a Pi assay with and without incubation with pyrophosphatase (PPA). We found that the majority of products accompanying ATP depletion by rTENT4B were PPi irrespective of oligonucleotide addition (**Fig. 1B**), and that a catalytically dead rTENT4B did not consume ATP (**Supplementary Fig. 1A**), arguing against a contaminating ATPase activity in the rTENT4B purification process that would be expected to generate Pi. We next asked whether contaminating RNAs in trace amounts in the rTENT4B preparation could be serving as primers to underlie the ATP consumption and PPi generation. To assess this, we treated 100 pmoles of rTENT4B (100x the amount of RNA oligo used for comparison) with proteinase K, purified the residual material under conditions for isolating RNA, and subjected it to 5’ end labeling with γ-^32^P-ATP and polynucleotide kinase. One pmole of a 15-mer homopolymeric adenosine RNA oligonucleotide (oligo A15) was also end-labeled and used as a control. However, we were unable to detect labeled RNA in this manner, suggesting that significant quantities of contaminating RNA species in the rTENT4B prep do not serve as substrates to account for the observation of primer-independent ATP consumption by rTENT4B (**Supplementary Fig. 1B**). These data indicated an unusual convergence of both primer- and template-independent PPi production from ATP by a polymerase.

**Fig. 1.**
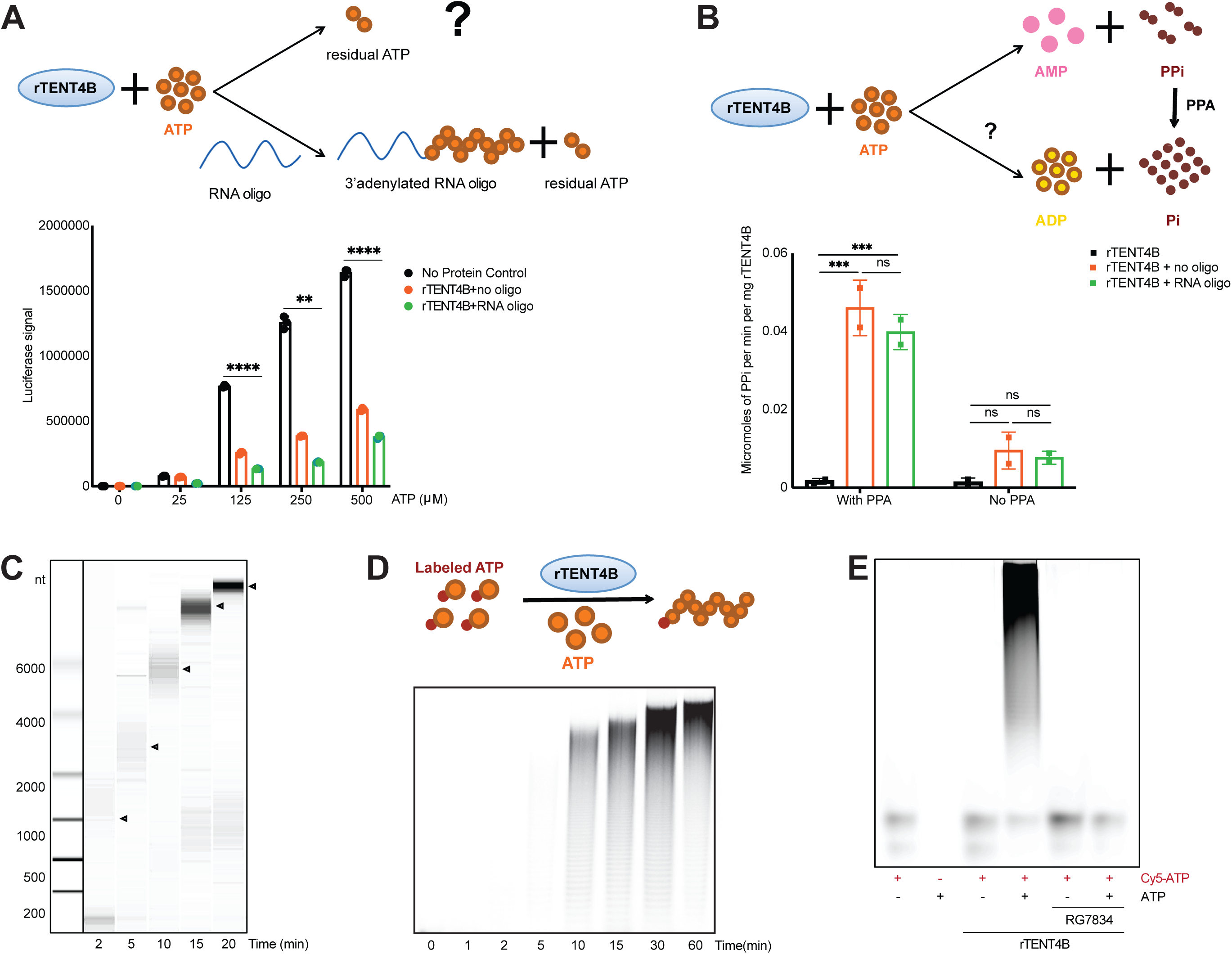
De novo ATP polymerization by rTENT4B leads to RNA synthesis and RNA-substrate 3’extension. **A,** *Upper:* schematics of ATP consumption by rTENT4B in the presence/ absence of an RNA-oligo as substrate and generation of residual ATP. *Lower:* luciferase-based detection of ATP consumption by rTENT4B in the presence of an RNA-substrate oligo compared to no RNA-substrate added reactions. ATP alone in the absence of rTENT4B was used as control (no protein control). n=3 biological replicates. Mean ± S.D. shown, 2-way ANOVA, **p<0.01, ****p<0.001. **B,** *Upper:* Schematics of ATP conversion and detection via pyrophosphate (PPi) assay, depending on the presence of inorganic pyrophosphatase (PPA), PPi or Pi would be released. *Lower:* Consumption of ATP by rTENT4B via analysis of PPi yield, in the presence/ absence of inorganic pyrophosphatase (PPA) as a measure of ATP hydrolyzation (ADP + Pi), compared to 3’ extension of an RNA-substrate oligo (PPi). n=2 biological replicates. Mean ± S.D. shown, 2-way ANOVA, ns- not significant, ***p<0.005. **C,** De novo ATP polymerization using rTENT4B. rTENT4B was incubated with fixed concentration of ATP (100 µM) and was subjected to RNA-substrate extension assay in the absence of an RNA-substrate, and generation of de novo synthesized nucleic acid products were detected using Bioanalyzer (Agilent picoChip analysis). **D,** *Upper:* Schema of replacing RNA-substrate oligo with 5’-Cy5-blocked ATP as an initiator nucleotide in RNA substrate extension assay. *Lower:* 5’-Cy5-labeled ATP was subjected to 3’ adenylation by rTENT4B, showing generation of Cy5-fluorescent RNA chains of varying length, elongating over time. n=3, one representative shown. **E,** Comparison of 3’ adenylation by rTENT4B using 5’-Cy5-labeled ATP in the presence/absence of 100µM RG7834 (rTENT4B inhibitor) treatment, showing generation of Cy5-fluorescent RNA chains of varying length, disappeared in the presence of RG7834. n=3, one representative shown.

We next asked whether it was possible that rTENT4B was mediating de novo nucleotide polymerization from free ATP molecules. To test this, we performed a time-course of rTENT4B reactions with varying concentrations of ATP and analyzed the products by polyacrylamide gel electrophoresis. In this manner, we detected concentration- and time-dependent increases in high molecular weight nucleic acid species, beginning at 10 µM ATP within 15 minutes (**Supplementary Fig. 1C**). Polymers >6000 nucleotides were detected within 15 minutes at higher ATP concentration (100 µM) indicating high processivity (**Fig. 1C**). ATP consumption in the luciferase assay correlated with the appearance of these products (**Supplementary Fig. 1D**). To definitively evaluate for de novo synthesis from a free ATP molecule, we next asked whether rTENT4B could initiate a polymer using a fluorescent γ-phosphate-labeled ATP plus unlabeled ATP. In these experiments, if γ-phosphate-labeled ATP is utilized for initiation of an RNA polymer, the label would be retained and thus detectable in progressively higher molecular weight species. Conversely, if a contaminating primer is initiating polymers rather than ATP, the γ-phosphate would be lost and labeled high molecular weight species would not be detectable. Remarkably, using γ-[(6-aminohexyl)-imido]-ATP-Cy5 (γ-Cy5-ATP) in the presence of rTENT4B, we were able to detect abundant fluorescently labeled polymers of increasing length as a function of unlabeled ATP concentration and time (**Fig. 1D, Supplementary Fig. 1E,F**). These data definitively demonstrate de novo primer- and template-independent polymerization of free ATP molecules by a eukaryotic polymerase, TENT4B.

We next verified that the core catalytic activity of rTENT4B is required for de novo polymerization of ATP. We have previously shown that pharmacological inhibition of rTENT4B prevents RNA oligo 3’ adenylation. Thus we tested RG7834, a potent TENT4B inhibitor discovered in HbSAg screens(30,41,42), for effects on its de novo nucleotide polymerization activity. We detected newly synthesized fluorescent chains with Cy5-labeled-ATP in the presence of rTENT4B, but not after RG7834 treatment (**Fig. 1E**). These data support de novo RNA synthesis as a catalytic property of rTENT4B that is amenable to pharmacological inhibition. Next, based on reports that a conserved motif of basic amino acids at the C-terminus of rTENT4B is required for RNA substrate binding (25), we tested the ability of a C-terminal truncation construct (rTENT4B-delC) to mediate primer-free RNA polymerization. Again, we found that fluorescent RNA chains could be formed by rTENT4B-delC using γ-Cy5-ATP as the initiating nucleotide (**Supplementary Fig. 1G**). Taken together, these results demonstrate that the core catalytic function of rTENT4B includes a capacity for primer- and template-independent polymerization of free ATP molecules. (**Supplementary Fig. 1H**).

### High and unique efficiency of *de novo* ATP polymerization by rTENT4B

We next interrogated the relative efficiency of de novo versus primer-dependent nucleotide polymerization by rTENT4B. To do this, we quantified PPi production from ATP by rTENT4B in the presence or absence of an RNA oligonucleotide. We found that the generation of PPi by rTENT4B increases in an ATP concentration-and time-dependent manner (**Fig. 2A,B**). However, PPi generation is not significantly different in the presence or absence of 25 pmoles oligonucleotide primer, at concentrations as low as 5 µM ATP and reaction durations from 30 minutes to plateau at 180 minutes (**Fig. 2A,B**). We next asked whether other RNA polymerases showed the capacity for de novo NTP polymerization. We began by testing canonical RNA PAPs and observed no yield of PPi when incubating either eukaryotic (yeast) PAP or prokaryotic (*E. coli*) PAP with 100 µM ATP in the absence of an RNA oligonucleotide (**Fig. 2C**). In contrast, we observed a significant increase in PPi yield with yeast PAP at 1000 µM ATP without an oligonucleotide, whereas *E.coli* PAP failed to yield PPi at 1000 µM ATP without an oligonucleotide (**Fig. 2D, Supplementary Fig. 2A,B**). No significant difference was observed with 1000 µM versus 100 µM ATP using rTENT4B, with or without an oligonucleotide (**Supplementary Fig. 2C)**. No Cy5-labeled high molecular weight polymers were observed with yeast PAP when tested using γ-Cy5-ATP and 1000 µM unlabeled ATP **(Fig. 2E).** However, high molecular weight fluorescent signals indicating nucleic acid polymers were detected using Sybr gold intercalating dye on the same denaturing gels (**Fig. 2E**). This result potentially indicates that yeast PAP cannot utilize γ-Cy5-ATP as an initiating NTP as opposed to ATP. Alternatively, we cannot rule out that the yeast PAP preparation might contain contaminating RNA that is acting as a primer. Taken together, these data indicate a highly efficient and potentially unique capacity for TENT4B to polymerize γ-Cy5-ATP and unlabeled ATP relative to canonical PAPs.

**Fig. 2.**
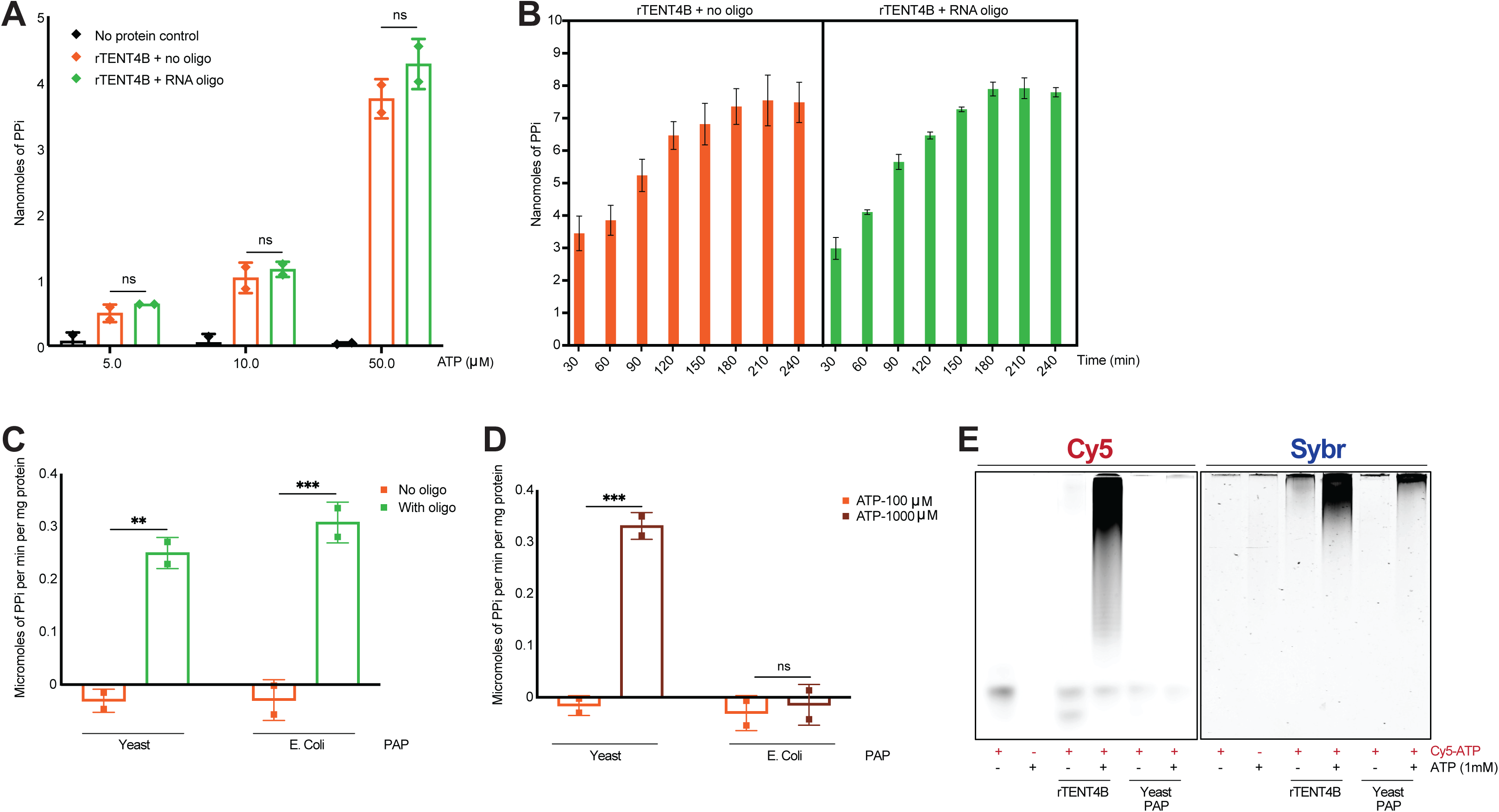
Rate of de novo ATP polymerization by rTENT4B compared to poly A-polymerases. **A,** Analysis of PPi generated by rTENT4B in response to increasing concentrations of ATP, in the presence of an RNA-oligo as substrate, compared to no RNA-substrate added reactions, and ATP alone in the absence of rTENT4B (no protein control). n=2 biological replicates. Mean ± S.D. shown, 2-way ANOVA, ns- not significant. **B,** Analysis of PPi yield generated by rTENT4B in the presence/ absence of an RNA-oligo as substrate as a measure of ATP consumption over time. n=2 biological replicates. Mean ± S.D. shown. **C,** Comparison of PPi yield generated by canonical PAPs from yeast and bacteria (*E. coli.* PAP) via using 100µM ATP as substrate in the presence/absence of an RNA oligo as substrate in 30 minutes. n=2 biological replicates. Mean ± S.D. shown, 2-way ANOVA, **p<0.01, ***p<0.005. **D,** Comparison of PPi yield generated by canonical PAPs from yeast and bacteria (*E. coli.* PAP) via using 100µM and 1000µM ATP as substrate in no-RNA oligo added reactions in 30 minutes. n=2 biological replicates. Mean ± S.D. shown, 2-way ANOVA, ns- not significant, ***p<0.005. **E,** Comparison of ATP polymerization using rTENT4B and canonical PAP from yeast using 5’-Cy5-labeled ATP as substrate along with 1000µM ATP. ATP incorporation was confirmed via detecting generation of Cy5-labeled RNA chains after 30 min in no-RNA oligo added reactions. Unlabeled nucleic acid products generated during the reaction were detected via staining with Sybr gold. n=3, one representative shown.

We next tested other noncanonical PAPs including rTENT4A, rTENT2 and CID1 for their ability to catalyze primer-independent de novo RNA synthesis. As expected, all recombinant enzymes could extend RNA oligonucleotide primers (**Supplementary Fig. 2D**). However, we observed limited high molecular weight fluorescent signals using Sybr gold in the absence of RNA oligonucleotides using these noncanonical PAPs plus 100-1000 µM ATP (**Supplementary Fig. 2D,E**). Taken together, we find that other noncanonical polymerases might also be able perform de novo synthesis of RNA polymers using ATP as a substrate, albeit less efficiently than rTENT4B.

### De novo, self-limiting polymerization by rTENT4B using GTP

TENT4B has been shown to utilize NTPs other than ATP in vitro and in cells (24,25,27,29). By differential scanning fluorimetry, we found that rTENT4B has the capacity to bind and thus be stabilized by any of the four canonical NTPs (**Supplementary Fig. 3A**). By primer extension using varying NTP concentrations followed by deep sequencing, we verified the capacity of rTENT4B to utilize any of these for RNA polymerization, recapitulating the preference ATP > GTP >> CTP / UTP (**Supplementary Fig. 3B-C**). We next sought to assess the promiscuity of NTP utilization by rTENT4B in de novo RNA synthesis. We first used γ-Cy5-ATP with unlabeled NTPs and analyzed the reaction products for synthesis of fluorescent RNA polymers. We found that rTENT4B could generate polymers starting with γ-Cy5-ATP at high concentrations of GTP (1 - 3.3 mM) but not CTP or UTP (**Fig. 3A**). Next, we tested de novo GTP polymerization using rTENT4B in the absence of γ-Cy5-ATP. Remarkably, we detected strong and discrete oligonucleotide products by Sybr gold staining at GTP concentrations (1.5 – 5 mM) similar to those observed in γ-Cy5-ATP experiments (**Fig. 3B**). We verified that the polymerization of free GTP nucleotides by rTENT4B was accompanied by PPi generation (**Fig. 3C**). Interestingly, the polymers created using GTP were restricted to ≤20 nt, versus >6000 nt for ATP at similar concentrations and time frames (**Fig. 1C, 3A,B**; *see below*). Next, we tested whether de novo chains could be initiated with γ-(6-Aminohexyl)-GTP-Cy5 (γ-Cy5-GTP) rather than γ-Cy5-ATP, using unlabeled ATP. We found robust formation of high molecular weight species retaining a 5’ Cy5 label when we added unlabeled ATP, that increased in a concentration-dependent manner (**Fig. 3D**). Finally, we analyzed the capacity of rTENT4B to utilize any NTP at high concentrations in the absence of an RNA primer and analyzed the results with Sybr gold staining. This experiment confirmed de novo synthesis of nucleic acid polymers using ATP or GTP but not CTP or UTP. Again, chain formation was self-limiting with GTP (**Fig. 3E**). These data demonstrate de novo polymerization of GTP by rTENT4B, in a self-limiting manner compared to ATP.

**Fig. 3.**
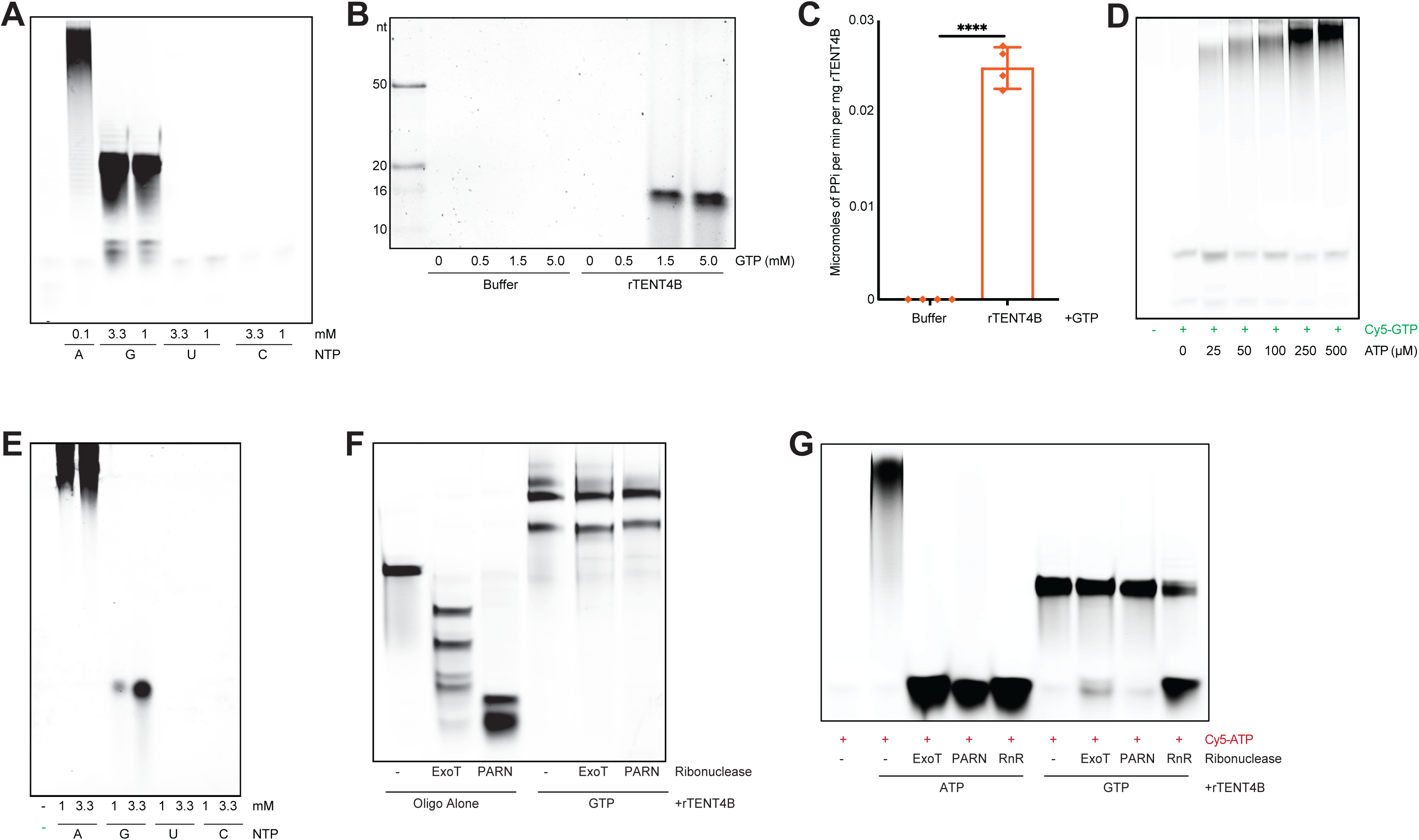
De novo NTP polymerization by rTENT4B. **A,** rTENT4B along with four canonical NTPs-ATP (A), CTP (C), UTP (U), GTP (G), plus 5’-Cy5-labeled ATP as substrate. NTP incorporation was confirmed via detecting generation of Cy5-labeled RNA chains after 30 min in no RNA-substrate added reactions. n=3, one representative shown. **B**, De novo GTP polymerization using rTENT4B. rTENT4B along with increasing concentrations of GTP was subjected to RNA extension assays in no-RNA substrate added reactions, and generation of de novo synthesized nucleic acid products were detected using denaturing TBE/Urea-acrylamide gel followed by Sybr gold staining. n=3, one representative shown. **C**, rTENT4B along with GTP was subjected to RNA-substrate extension assay in the absence of an RNA-substrate, and GTP polymerization was detected via PPi analysis. n=4 biological replicates as shown. Mean ± S.D., two-tailed unpaired t test ****p<0.001. **D,** 5’-Cy5-labeled GTP was subjected to 3’ adenylation using increasing concentrations of ATP and rTENT4B, showing generation of fluorescent (Cy5)-labeled RNA chains of varying length. n=3, one representative shown. **E,** De novo NTP polymerization using rTENT4B. rTENT4B along with increasing concentrations of four canonical NTPs was subjected to RNA-substrate extension assay in no RNA-substrate added reactions, and generation of de novo synthesized RNA products were detected using denaturing TBE/Urea-acrylamide gel followed by Sybr gold staining. n=3, one representative shown. **F,** 3’ exonuclease degradation assay using exonuclease T (ExoT) or recombinant purified PARN (PARN), for unmodified FAM-labeled oligo 3 (Oligo Alone) versus rTENT4B-mediated GTP-tailed oligo. n=3, one representative shown. **G,** 3’ exonuclease degradation assay using exonuclease T (ExoT) or recombinant purified PARN (PARN) or single strand specific RNase R (RnR), for de novo synthesized Cy5-fluorescent RNA products by rTENT4B, in no RNA-substrate added reactions, via using Cy5-labeled ATP alone versus Cy5-labeled ATP plus unlabeled-ATP or unlabeled-GTP. n=2, one representative shown.

Earlier we have demonstrated that rTENT4B can utilize 2’-O-methyl-ATP to form a self-limiting polymer on RNA 3’ ends, in a manner that both constrained further nucleotide addition and also conferred 3’ exonuclease resistance(48). We therefore hypothesized that the self-limiting nature of oligo-G polymer formation might reflect formation of a secondary structure that is resistant not only to further enzymatic polymerization but also enzymatic degradation. In keeping with this, we found that an RNA oligonucleotide extended using rTENT4B and GTP formed discrete high molecular weight species that unlike the unmodified oligonucleotide were completely resistant to 3’ exonucleases, exonuclease T (exoT) and recombinant PARN (rPARN) (**Fig 3F**). Next, we tested whether the de novo synthesized RNA products produced by rTENT4B from γ-Cy5-ATP plus unlabeled ATP versus unlabeled GTP were differentially sensitive to exonucleases. Indeed, we found that γ-Cy5-labeled polymers formed by rTENT4B using GTP were resistant to exoribonucleases compared to those generated via using ATP (**Fig 3G**). Ribonuclease R (RnR) is a 3’ exonuclease with a characteristic capacity to degrade highly structured RNAs. We found that GTP-formed oligonucleotides were sensitive to RnR compared to exoT and rPARN (**Fig 3G**), in keeping with the notion that a secondary structure underlies their nuclease-resistance. Collectively, these results demonstrate the ability of rTENT4B to utilize not only free ATP but also GTP in de novo RNA synthesis, but in a self-limiting manner yielding exonuclease resistant polymers.

### Purine nucleotide diphosphates serve as substrates for rTENT4B

We have previously shown that rTENT4B could utilize adenosine diphosphate (ADP) in RNA oligonucleotide extension reactions (**Fig. 4A**)(48). We thus asked whether other nucleoside diphosphates (NDPs) could serve as substrates for rTENT4B in RNA polymerization. In a primer-extension assay, we first determined rTENT4B could utilize GDP but not UDP or CDP, and only at higher concentrations than ADP (1 mM versus 0.1 mM) (**Fig. 4B**). As with GTP, short, self-limiting polymers were observed with GDP compared to those generated using ADP (**Fig. 4B**). We next tested de novo RNA synthesis via using NDPs and rTENT4B in primer-free reactions using γ-Cy5-ATP as a substrate and observed robust γ-Cy5 polymer formation using ADP and GDP. As expected, no fluorescent polymers were generated by rTENT4B when incubated with CDP or UDP (**Fig. 4C**). When free ADP was utilized as a substrate for rTENT4B without an RNA primer or γ-Cy5-ATP, high molecular weight RNA polymers could be detected using Sybr gold within 1 hour, albeit at high concentrations ≥ 3.3 mM (**Fig. 4D**). When using free ADP as a substrate, Pi was generated instead of PPi, as expected (**Fig. 4E**). While we cannot rule out the possibility of trace ATP contamination serving an initiating nucleotide at these high concentrations of ADP, our data collectively suggest that rTENT4B can utilize purine diphosphate nucleotides in primer- and template-independent RNA synthesis.

**Fig. 4.**
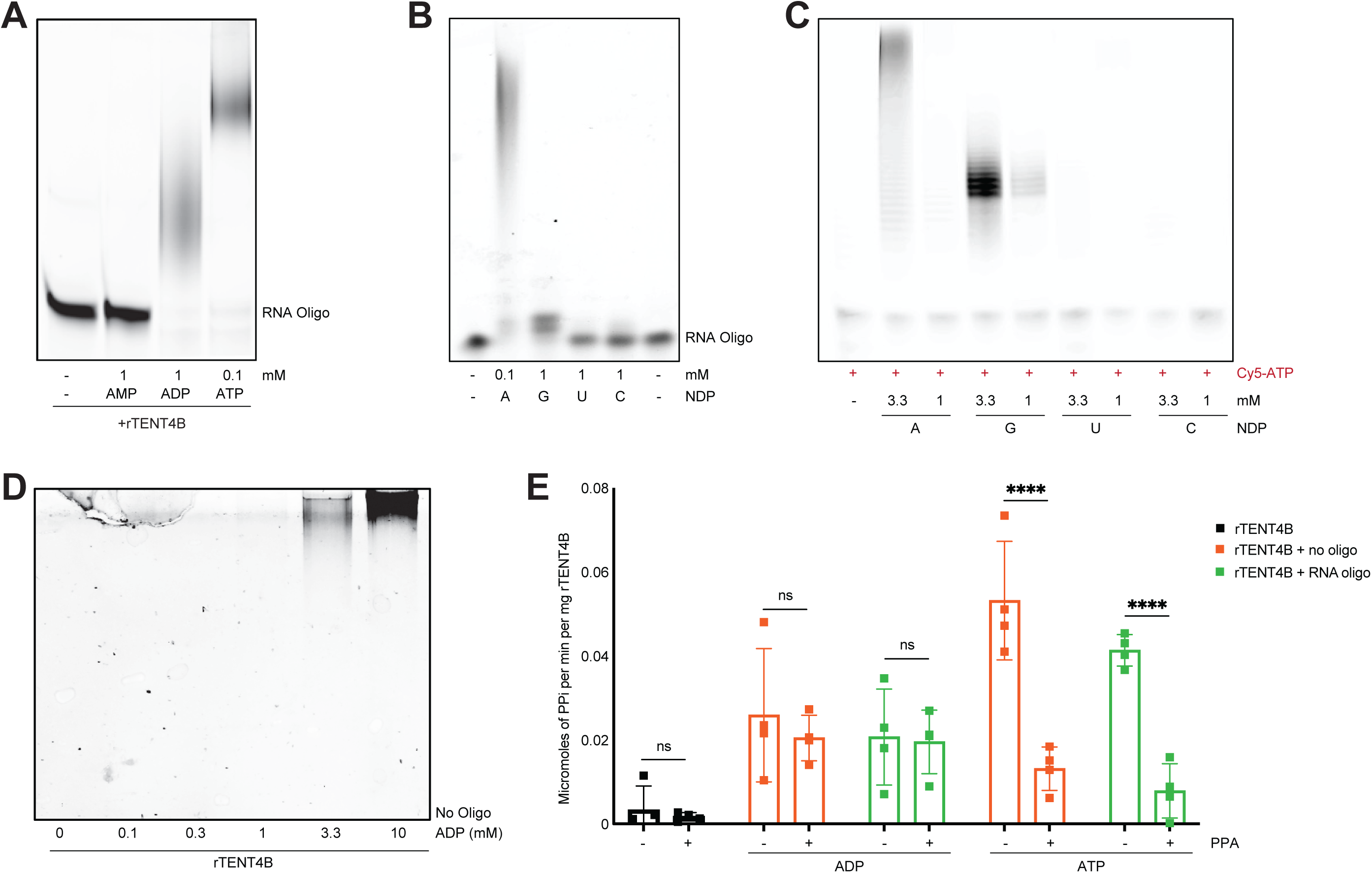
Purine nucleotide diphosphates (NDPs) polymerization by rTENT4B. **A,** Extension of a 20-mer RNA substrate oligo (oligo 1) using rTENT4B and ATP analogs demonstrating differential incorporation. (AMP: Adenosine-5’-monophosphate; ADP: Adenosine-5’-diphosphate; ATP: Adenosine-5’-triphosphate. n=3 replicates, one representative shown. **B,** Utilization of nucleotide diphosphates (NDPs) by rTENT4B. rTENT4B along with a 20-mer RNA-substrate oligo (oligo 2) was incubated in the presence of four canonical NDPs, in RNA-substrate extension assays, and reaction products were resolved using denaturing TBE/Urea-acrylamide gel followed by Sybr gold staining. **C**, rTENT4B was incubated with four canonical NDPs plus 5’-Cy5-labeled ATP as substrate. NDP incorporation was confirmed via detecting generation of Cy5-labeled RNA chains after 30 min in no RNA-substrate added reactions. **D**, De novo ADP polymerization using rTENT4B. rTENT4B along with increasing concentrations of ADP was subjected to RNA-substrate extension assay in the absence of an RNA-substrate, and generation of de novo synthesized RNA products were detected using denaturing TBE/Urea-acrylamide gel followed by Sybr gold staining. **E**, Consumption of ADP by rTENT4B via analysis of Pi yield, in the presence/ absence of inorganic pyrophosphatase (PPA) as a measure of ATP contamination (ADP + Pi), in the presence/absence of an RNA-substrate oligo, compared to that of rTENT4B alone versus rTENT4B plus ATP. n=4 biological replicates. Mean ± S.D. shown, 2-way ANOVA, ns- not significant, ****p<0.001.

## Discussion

Our data show primer- and template-free polymerization of RNA from free nucleotides by TENT4B, a property that has not been definitively ascribed to a polymerase in the past. Qß replicase is an RNA-dependent RNA polymerase from E. coli bacteriophage Qß previously proposed to be capable of catalyzing de novo synthesis of various RNA oligonucleotides from free NTPs, i.e. in a primer- and template-independent manner(49). While initial concerns about potential RNA primer contamination(50) were addressed using more rigorous purification methods(51), it was ultimately found that Qß replicases utilize short host cell-derived RNA templates in a reaction that becomes autocatalytic(52,53). Thus Qß replicase conforms to classification as a primer-independent but template-directed RNA polymerase. In 1955, polynucleotide phosphorylase (PNPase) from *Azotobacter vinelandii* was shown to polymerize RNA from NDPs in vitro, enabling synthesis of polyribonucleotides used to crack the genetic code(54,55). While PNPase participates in RNA synthesis in prokaryotic cells under certain circumstances(56), the primary role of PNPase across diverse species is RNA degradation via phosphorolysis(57). To our knowledge, this report is the first of primer- and template-free RNA synthesis by a eukaryotic polymerase.

Our experimental demonstration of de novo RNA polymerization using rTENT4B exploited its ability to utilize γ-Cy5-ATP or γ-Cy5-GTP as the initiating nucleotide. In retaining the 5’ label, single nucleotide additions were readily visualized using free purine nucleotides. We did not detect a lag in reaction kinetics as measured by PPi production without an oligonucleotide, indicating that engagement of a free nucleotide by rTENT4B in the “product” (P)-site of the polymerase active appears to be as efficient as that of an RNA primer. This is further indicated by robust activity of a C-terminal mutant of rTENT4B that does not contain a putative RNA binding site. Once initiated, peak PPi generation and extent of product formation were similar with or without an RNA oligonucleotide primer. These data and the 1000s-mer reaction products rapidly generated using ATP indicate that rTENT4B-mediated RNA polymerization is highly processive in vitro and limited only by the available ATP pool.

The physiological relevance of the novel properties and capabilities of rTENT4B we have characterized here is not known. TENT4B is the human ortholog of yeast Trf4p, which forms the TRAMP complex with Air2 and Mtr4p involved in RNA quality control (58-61). Trf4p oligo-adenylates nascent nuclear RNAs that are potentially misfolded with an available 3’ end. Via the TRAMP complex, these modified RNAs are targeted for rapid degradation by the nuclear RNA exosome. Similarly, human TENT4B forms multi-protein complexes with orthologs of Air2 and Mtr4, ZCCHC7 and MTR4 respectively(62,63), catalyzing oligoadenylation-mediated exosome-targeting of incomplete and cryptic promoter upstream transcripts and other RNAs including TERC and snoRNAs(46,64-66). In vitro, yeast Mtr4p limits poly-adenylation by Trf4p by stimulating its dissociation from substrate after addition of 3-4 adenosines(67). In human cells, RNA targets such as TERC and scaRNA13 display oligo-adenylation (<10 nt) rather than poly-adenylation when TENT4B activity is unopposed due to PARN disruption(68), also suggesting that TENT4B processivity is more constrained in a physiological setting than what we have observed in vitro. However, a recent report implicates TENT4B in untemplated 3’ tailing of rRNA fragments of several hundred nucleotides long (so-called starRNA) in the setting of amyloidogenesis(69), suggesting that under certain conditions TENT4B may display higher processivity in cells. We have previously demonstrated that complete inactivation of TENT4B by CRISPR/Cas9 is tolerated in human cells without adverse effects on cell viability or growth(24,70). Nevertheless, a role for de novo or processive NTP polymerization by TENT4B in vivo remains to be determined.

Despite showing a preference for purines, rTENT4B was capable of condensing all four canonical nucleotides into polymers, an observation with potential relevance in biotechnology. Complete chemical synthesis of short RNA species has enabled novel therapeutics such as anti-sense oligonucleotides (ASOs) and short interfering RNAs (siRNAs)(71) but has limitations in terms of polymer length and synthetic capacity, including environmental concerns at large scale due to use of hazardous chemicals(72). Recent studies demonstrate enzymatic synthesis of custom short RNA polymers using mutant forms of CID1 and reversible terminator NTPs, but this approach relied on a pre-existing oligonucleotide of at least 10 nt in length followed by an endonucleolytic cleavage step(73). Given TENT4B’s homology to CID1 as members of the superfamily of polymerase beta-like nucleotidyltransferases(74), and its inherent NTP promiscuity relative to other nucleotidyltransferases for canonical and modified ribonucleotides(25,48), it is tempting to speculate that engineered forms of TENT4B could be used for de novo, template-independent enzymatic synthesis of RNAs in aqueous solution, without relying on an initiating oligonucleotide. As such, further studies of rTENT4B could advance the emerging field of complete enzymatic RNA synthesis, which promises to overcome barriers including polymer length and composition, scale, and environmental safety that limit the expansion of RNA biotechnology and therapeutics.

## Data Availability

The data that support the findings of this study are available within the manuscript and associated Supplementary data files. Unprocessed data will be provided on request. There are no restrictions on data availability from this study.

## Acknowledgements

N.N. discloses support for the research described in this study from BCH Department of Pediatrics and Manton Center for Orphan Disease Research, Uplifting Athletes Young Investigator Draft, and Team Telomere. Y.W.F. discloses support for the research described in this study from NIH (R01GM152463). A.K.T. discloses support for the NGS work described in this study from the Tufts University Core Facility Genomics Core (RRID:SCR_012501). S.A. discloses support for the research described in this study from NIH (R01DK107716) and the BCH Translational Research Program. This content is solely the responsibility of the authors and does not necessarily represent the official views of the National Institutes of Health.

## Author contributions

N.N. and S.A. conceived this study and designed the experiments. N.N. performed *in vitro* and biochemical analyses. A.K.T. performed computational analyses. Y.W.F. purified recombinant proteins. S.A. and N.N. wrote the manuscript. All authors edited the manuscript.

## Conflict of interest statement

N.N and S.A. are listed as co-inventors on patent applications including the work described in this paper.

## Material availability

Request for materials, reagents and resources used in this study should be directed to the corresponding author, Suneet Agarwal. Certain materials requests will require a completed Materials Transfer Agreement.

---

**Supplementary Fig. 1.**
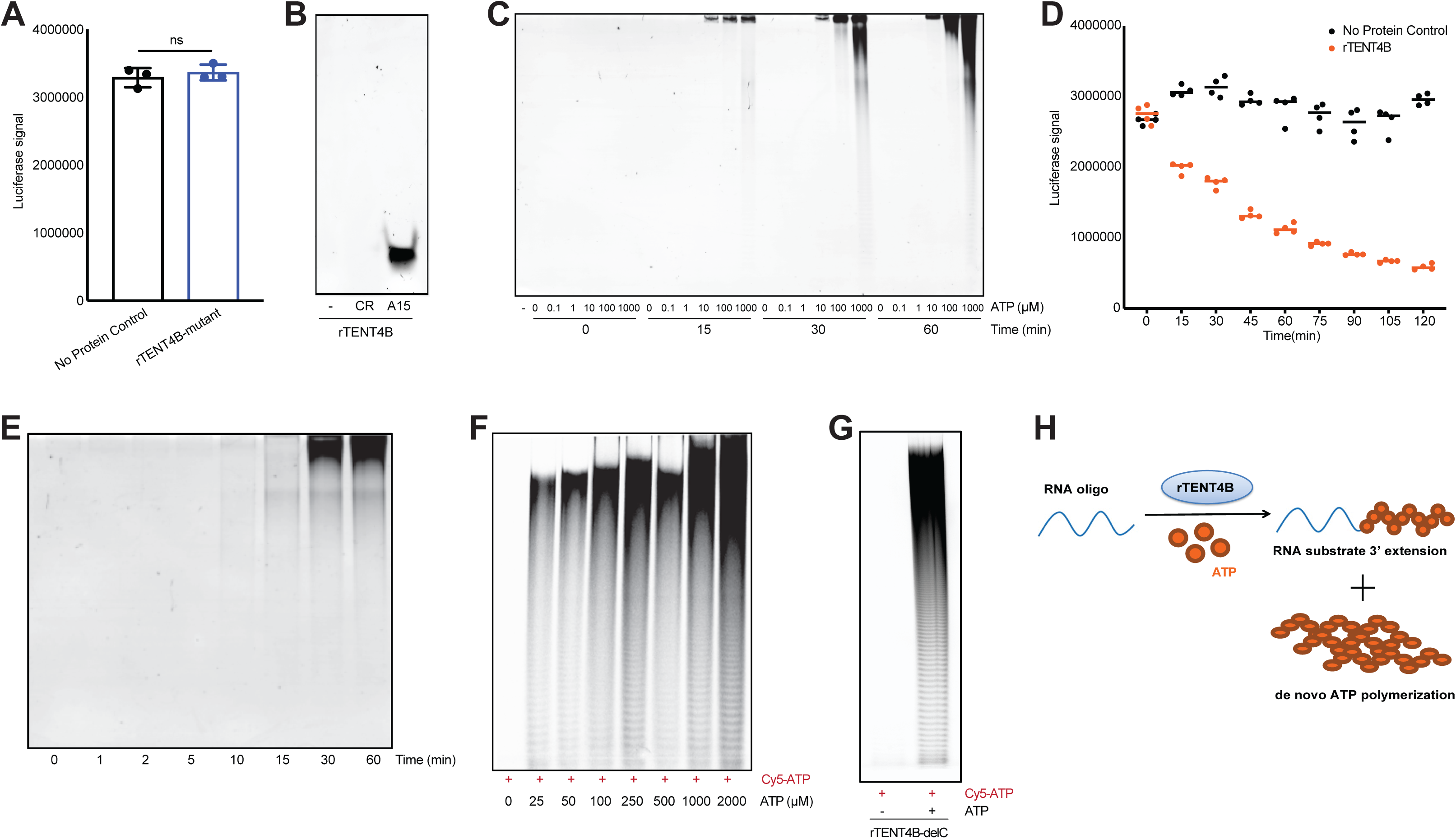
ATP polymerization by rTENT4B in the absence of an RNA-substrate. **A,** Luciferase-based detection of ATP (500 µM) consumption by catalytic site mutant rTENT4B in no RNA-substrate added reactions. ATP alone in the absence of rTENT4B was used as control (no protein control). n=3 biological replicates. Mean ± S.D. shown, two-tailed unpaired t test, ns- not significant. **B,** 100 pmoles of rTENT4B (CR or contaminating RNA) and one pmole of an unlabeled RNA-oligo as substrate (homopolymer of A, oligo A15) were subjected to proteinase K treatment and reactions were purified. Purified reaction products were then radio-labeled using 100 µM of α-p32 using rTENT4B. Reaction products were compared via autoradiography on a denaturing TBE/urea acrylamide gel to see the contaminating RNA (CR) species that may have purified while generating recombinant TENT4B. **C,** De novo ATP polymerization using rTENT4B. rTENT4B along with increasing concentrations of ATP was subjected to RNA-substrate extension assay in the absence of an RNA-substrate, and generation of de novo synthesized RNA products were detected over time using denaturing TBE/urea acrylamide gel followed by Sybr gold staining. **D,** ATP consumption over time by rTENT4B compared to no protein reaction mix as control, read out as luciferase signal in a luciferase-based assay. n=4 biological replicates, Mean ± S.D. shown. **E,** Visualization of non-fluorescent nucleic acid products generated via de novo ATP polymerization using Sybr gold staining, 5’-Cy5-labeled ATP was subjected to 3’ adenylation by rTENT4B, showing generation of fluorescent-labeled RNA chains of varying lengths, elongating over time. n=3, one representative shown. **F,** 5’-Cy5-labeled ATP was subjected to 3’ adenylation by rTENT4B, showing generation of fluorescent-labeled RNA chains of varying length with increasing concentrations of unlabeled ATP. n=3, one representative shown. **G,** C-terminal deletion mutant of rTENT4B (delC-TENT4B) was incubated with 250 µM of ATP plus 5’-Cy5-labeled ATP (100 µM), and generation of fluorescent-labeled RNA chains of varying lengths were detected using Cy5 channel. n=3, one representative shown. **H,** Schematics of dual ATP polymerization by rTENT4B to generate RNA-substrate 3’adenylation and ATP polyadenylation products.

**Supplementary Fig. 2.**
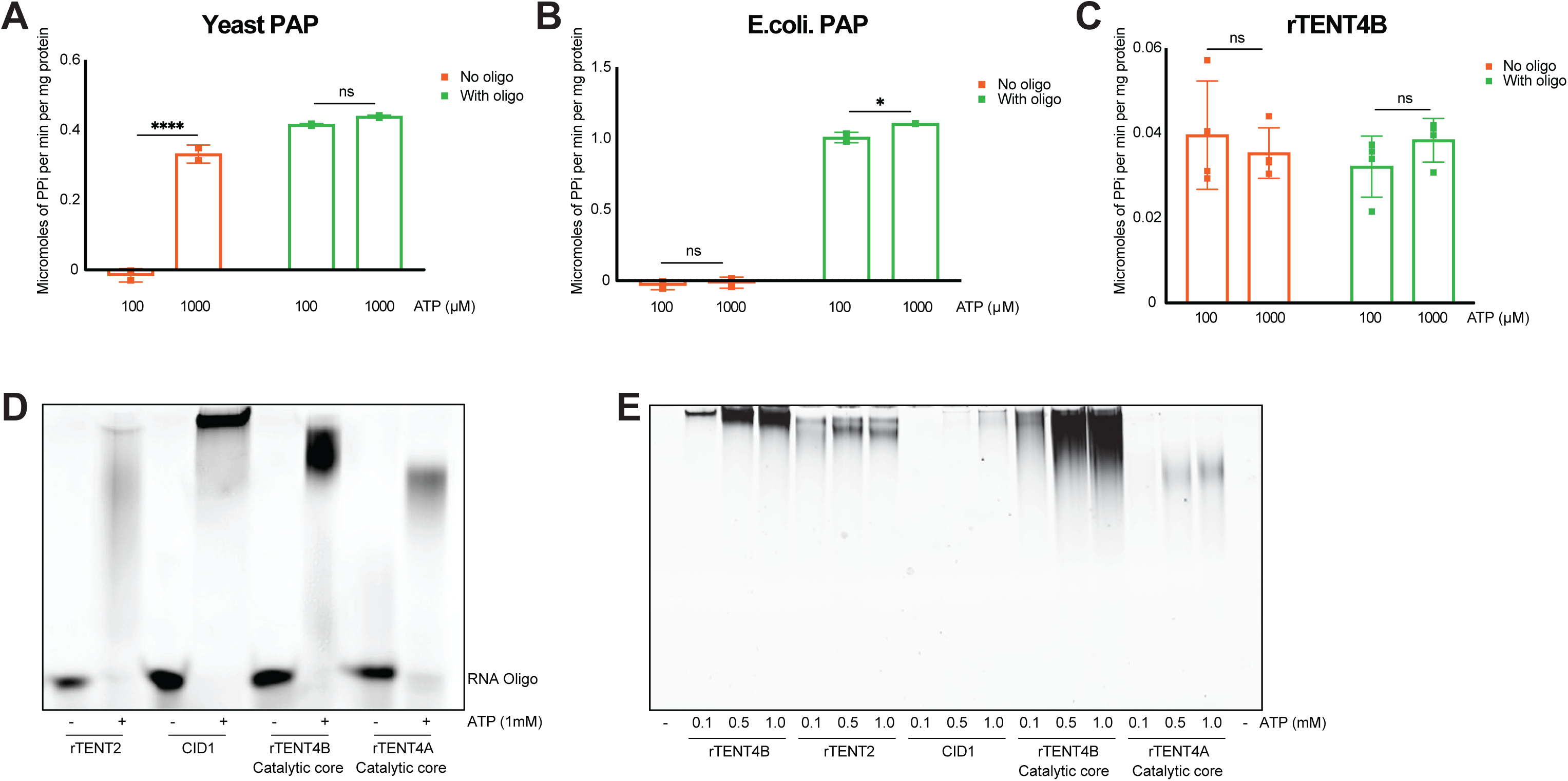
De novo ATP polymerization by rTENT4B and other eukaryotic poly(A)-polymerases. **A-C** Analysis of PPi generated by canonical PAPs from yeast (**A**) and bacteria (*E. coli.*) (**B**), compared to rTENT4B (**C**) via using 100 µM and 1000 µM ATP as substrate in no-RNA substrate added reactions compared to that of RNA-oligo added reactions. n=2 (n=4 for rTENT4B) biological replicates. Mean ± S.D. shown, 2-way ANOVA, ns- not significant, *p<0.05, ****p<0.001. **D,** RNA substrate extension by noncanonical PAPs or TENTs (or their catalytic cores) via using 1000µM ATP as substrate in the presence of a 20-mer RNA oligo (oligo 1). n=3, one representative shown. **E,** De novo ATP polymerization. Recombinant TENTs (or their catalytic cores) were incubated with increasing concentrations of ATP in the absence of an RNA-substrate, and generation of de novo synthesized RNA products were detected using denaturing TBE/urea acrylamide gel followed by Sybr gold staining. n=3, one representative shown.

**Supplementary Fig. 3.**
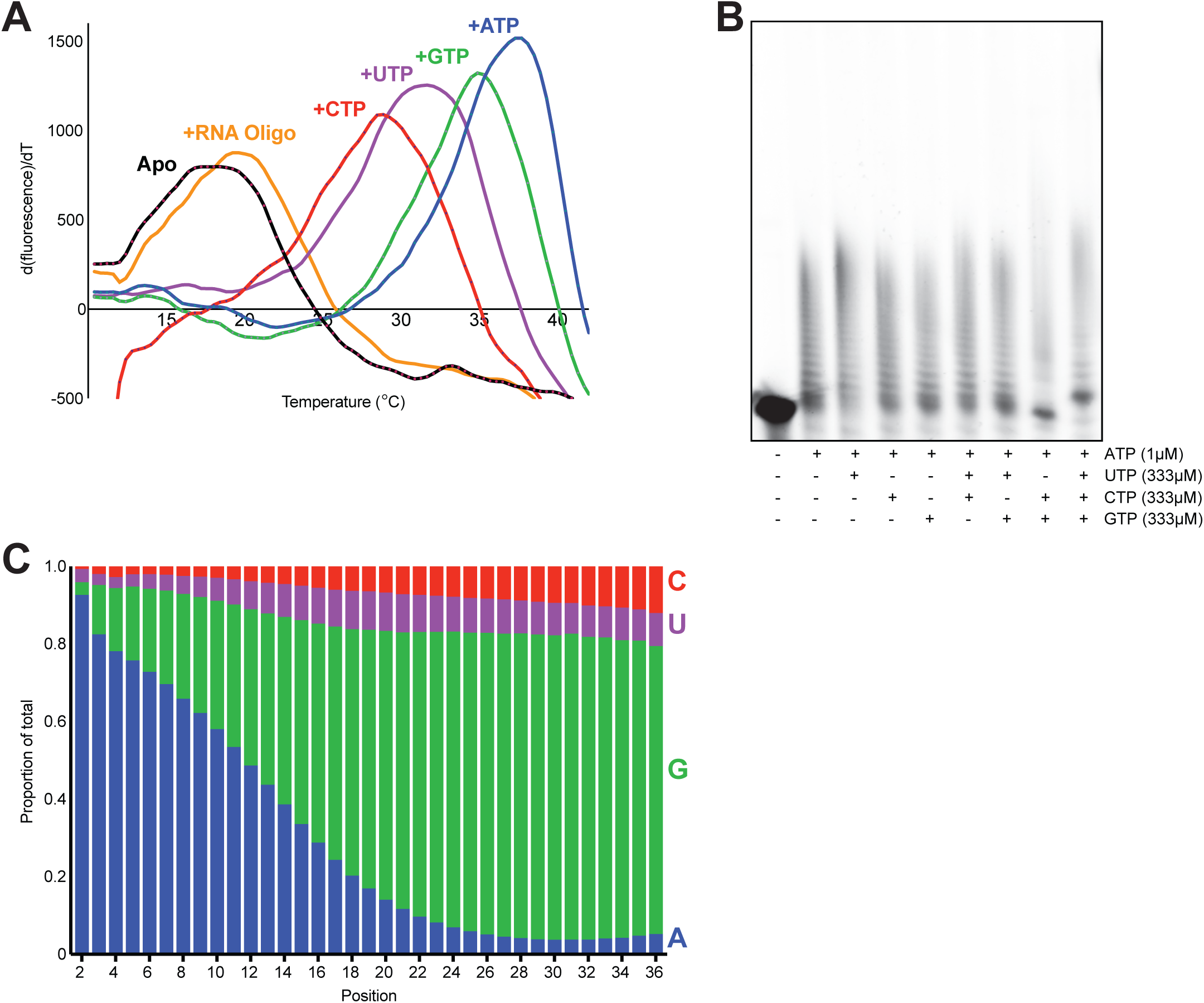
De novo NTP polymerization by rTENT4B prefers purines. **A,** Direct binding of an unlabeled RNA-substrate oligo (Oligo A15) or four canonical NTPs to rTENT4B, compared to protein alone (rTENT4B), by differential scanning fluorimetry (oligo- 1 pmole, NTPs- 100 µM each). **B-C,** Generation of homo-/hetero-polynucleotide chains by rTENT4B. rTENT4B along with RNA substrate oligo (Oligo 1) was incubated in the presence of four NTPs at indicated concentrations in figure, in RNA-substrate extension assays. Reaction products were resolved using denaturing TBE/urea acrylamide gel to visualize 3’ oligo extensions (**B**). Extended products (**B**) were subjected to deep sequencing after linker ligation followed by library preparation and analysis. The graph shows proportions (cut-off- 95% of total) for each base vs corresponding extended position from the oligo 3’end (**C**). n=2, one representative shown.

## Notes

### Competing Interest Statement

N.N and S.A. are listed as co-inventors on patent applications including work described in this paper.

